# Genomewide m^6^A mapping uncovers dynamic changes in the m^6^A epitranscriptome of cisplatin-treated apoptotic HeLa cells

**DOI:** 10.1101/2022.02.18.481057

**Authors:** Azime Akçaöz, Özge Tüncel, Ayşe Bengisu Gelmez, Buket Sağlam, İpek Erdoğan Vatansever, Bünyamin Akgül

## Abstract

Cisplatin, which is a traditional cancer therapeutic drug, induces apoptosis by modulating a diverse array of gene regulatory mechanisms. However, cisplatin-mediated changes in the m^6^A methylome is unknown. We employed m^6^A miCLIP-seq to investigate the effect of m^6^A methylation events under cisplatin-mediated apoptotic conditions in HeLa cells. Our high-resolution approach revealed numerous m^6^A marks on 972 target mRNAs with an enrichment on 132 apoptotic mRNAs. Following validation of site-specific m^6^A events on candidate RNAs, we tracked the fate of candidate mRNAs under *METTL3* knockdown and cisplatin treatment conditions. We detected perturbations in the translational efficiency of *PMAIP1* and *PHLDA1* transcripts based on the polysome profile analyses. Congruently, PMAIP1 and p53 amounts were dependent on METTL3. Additionally, cisplatin-mediated apoptosis was sensitized by *METTL3* knockdown. These results suggest that apoptotic pathways are modulated by m^6^A methylation events and METTL3-p53-PMAIP1 axis modulates cisplatin-mediated apoptosis in HeLa cells.

## Introduction

Apoptosis is a type of programmed cell death that is required to maintain the delicate balance between survival and cell death as part of cell and tissue homeostasis. Apoptosis, which is characterized by cytoplasmic shrinkage, chromatin condensation, nuclear fragmentation, and apoptotic bodies, is highly useful in eliminating damaged or unneeded cells without generating any inflammatory response (1). Thus, apoptotic processes are targeted by various chemotherapeutics drugs to treat cancer. For example, cisplatin induces apoptosis in cancer cells by inducing DNA lesions through intra- and inter-strand crosslinks in DNA (2,3). DNA lesions interfere with proper DNA replication and transcription, leading to the activation of DNA-damage response (DDR) and P53. P53 then transcriptionally coordinates the expression of genes that primarily triggers the intrinsic pathway of apoptosis as well as the extrinsic pathway (4). However, the efficacy of drugs plummets dramatically due to the drug resistance, which constitutes a major challenge in clinics for the proper treatment of cancer patients. Thus, it is imperative to understand the molecular mechanism of cisplatin-induced apoptosis to develop better strategies against drug resistance.

In an analogy to DNA and proteins, the chemical composition of RNA is modified co-or post-transcriptionally through epitranscriptomics mechanisms that play a vital role in the fate of modified RNAs (5). Of the existing 170 different types of epitranscriptomics modifications, the *N*^6^-methyladenosine (m^6^A) modification is the most abundant one with presence in 0.1-0.4% of all adenosines in cellular RNAs (5). Antibody-based enrichment coupled with high-throughput sequencing has uncovered m^6^A-methylated peaks not only on abundant tRNAs and rRNAs but also on mRNAs (6,7). Cell- and tissue-specific m^6^A modifications on mRNAs modulate a diverse array of cellular processes in health and disease. The emerging evidence suggests that the proteins and enzymes that orchestrate the cellular m^6^A dynamics control apoptosis under various cellular settings. For example, depletion of *METTL3* leads to an increase in the rate of apoptosis by reducing translation of MYC, BCL2 and PTEN in leukemia cells or BCL2 in breast cancer cells (8,9). The demethylase FTO, on the other hand, suppresses the rate of apoptosis in human acute myeloid leukemia cells (10). There are also reports on the modulatory effect of m^6^A reader proteins, such as IGF2BP1, on apoptosis (11). However, a complete m^6^A methylome of mRNAs under apoptotic conditions is not reported yet.

We used cisplatin, a universal cancer chemotherapeutic drug, as an inducer of apoptosis to profile m^6^A mRNA methylome in HeLa cells. Our analyses identified differential m^6^A methylation of 972 mRNAs. Interestingly, 132 of m^6^A-methylated mRNAs are associated with apoptosis as revealed by Gene Ontology analyses. Further analyses showed that cisplatin-induced m^6^A marks do not change the abundance of candidate mRNAs tested. However, we detected changes in the translational efficiencies of differentially methylated candidate mRNAs as revealed by polysome profiling. This observation was further supported by a corresponding increase in the protein amount, suggesting a potential role for METTL3-P53-PMAIP1 axis in apoptosis.

## Materials and Methods

### Cell Culture, Drug Treatments and Analysis of Apoptosis

HeLa cells, purchased from DKFZ GmbH (Germany), were cultured in RPMI 1640 (with 2 mM L-Glutamine, Gibco, USA) supplemented with 10% fetal bovine serum (FBS) (Gibco, United States) in a humidified atmosphere of 5% CO_2_ at 37°C. Cisplatin (Santa Cruz Biotechnology, United States) treatments were carried out in triplicates essentially as described previously (12). 0.1% DMSO was used as negative control for cisplatin. Treated cells were trypsinized by 1X Trypsin-EDTA (Gibco, United States) and washed in 1X cold PBS (Gibco, United States). Cell pellets were stained with Annexin V-PE (Becton Dickinson, United States) and 7AAD (Becton Dickinson, United States) in the presence of 1X Annexin binding buffer (Becton Dickinson, United States) followed by incubation at room temperature for 15 minutes in the dark. The rate of Annexin V and/or 7AAD-positive cells were quantified by a FACSCanto flow cytometer (Becton Dickinson, United States).

### Cell Transfection

0.6×10^6^ HeLa cells were seeded on 10 cm dishes (Sarstedt, Germany) and incubated overnight prior to transfection with 25 nM non-target pool (si-NC) or METTLl3 siRNA (si-METTL3) (Dharmacon, United States). DharmaFECT transfection reagent was mixed with siRNAs in a ratio of 2:1 (v/v) in 800-μl serum free medium and incubated for 5 minutes at room temperature. The DharmaFECT solution was added into the siRNA tube and incubated for 20 minutes at room temperature. The mixture was then combined with 8-ml medium prior to gently dropping over cells.

### Total RNA Isolation and qPCR

Total RNA isolation was performed using TRIzol™ reagent (Invitrogen, United States) according to the manufacturer’s protocol. Trace DNA contamination was eliminated by treating RNAs with TURBO DNA-free™ kit (ThermoFisher Scientific, United States) according to the manufacturer’s instructions.

For qPCR analyses, reverse transcription was conducted with RevertAid first strand cDNA synthesis kit according to the manufacturer’s instructions (Thermo Fisher Scientific, United States). cDNA was prepared using 2 μg of total RNA and diluted 1:10 in nuclease-free water. qPCR reactions were set up in triplicates with GoTaq® qPCR Master Mix (Promega, United States) and run in a Rotor-Gene Q machine (Qiagen, Germany). The primer sequences are listed in Table 1. All reactions were incubated at 95°C for 2 minutes as initial denaturation, 45 cycles of denaturation at 95°C for 15 seconds and annealing at 60°C for 1 minute following a melting step. GAPDH was used to normalize the qPCR data.

**Table 1.**
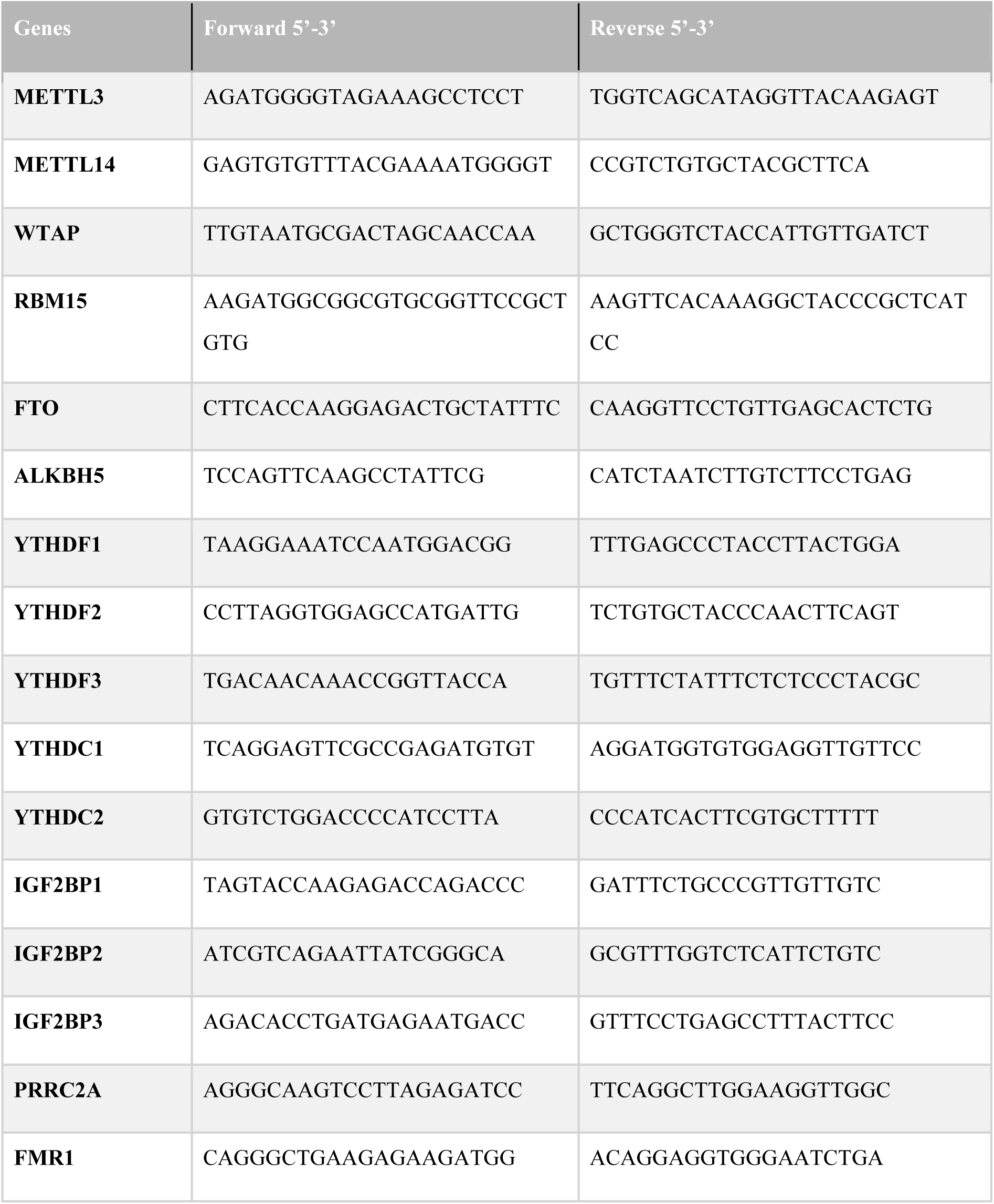
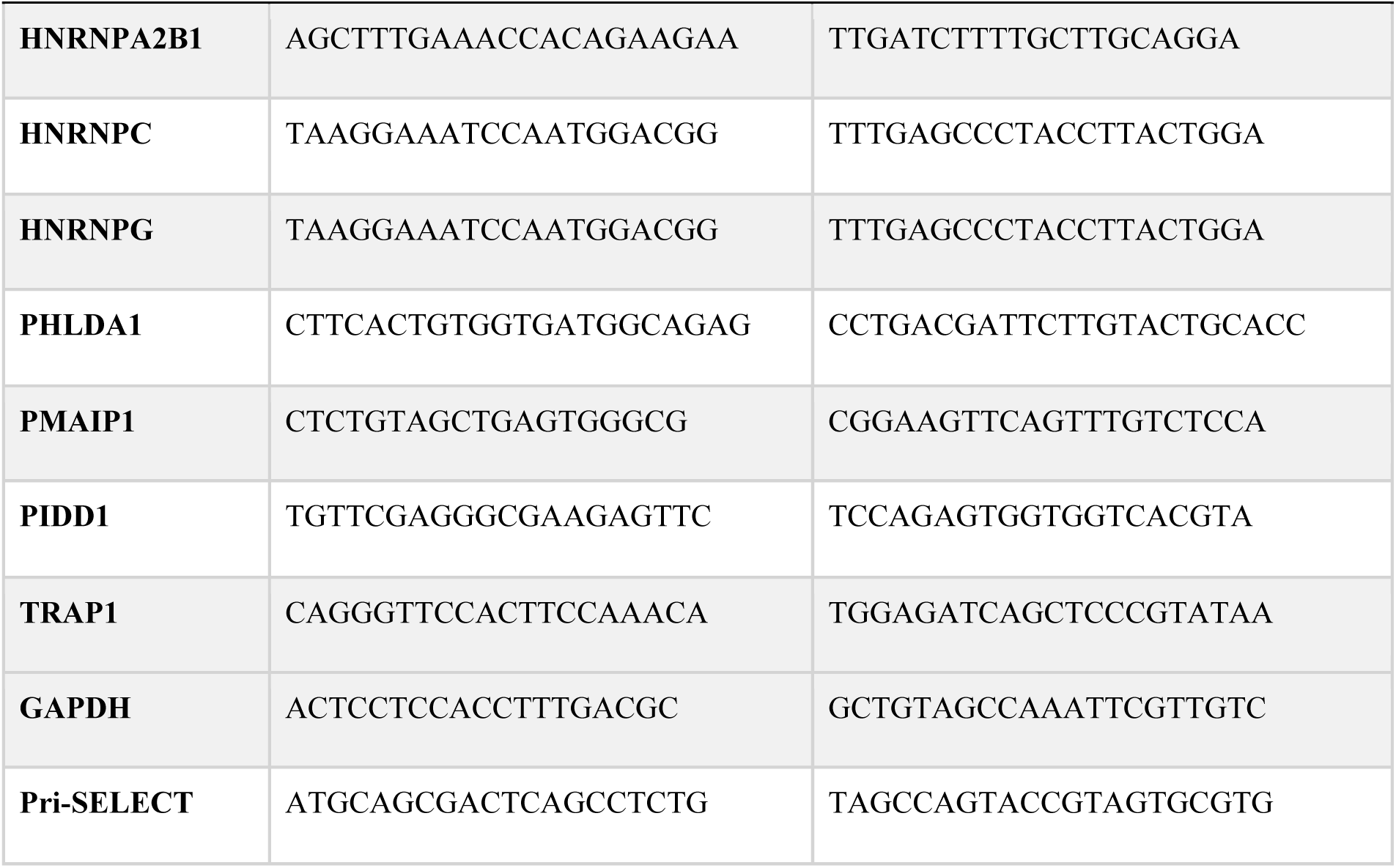
The list of primers used in qPCR analyses.

### m6A-miCLIP-seq and Bioinformatic Analysis

Three biological replicates of total RNAs isolated from DMSO-(0.1%) and cisplatin-treated HeLa cells were subjected to m^6^A-miCLIP-seq by Eclipse Bioinnovations (San Diego, USA) according to the published protocols (Linder et al., 2015). The data was deposited to GEO with the accession number GSE188580.

Data analysis was also performed by Eclipse Bioinnovations using their standard pipeline (13). Briefly, UMI sequences (the first 10 bases) were demultiplexed from each read by UMI-tools (14). Quality control of reads and adaptor trimming from 3’ ends was assessed by FastQC: v. 0.11.5 and Cutadapt: v. 1.15 tools, respectively. After the removal of contaminating sequences (e.g. rRNA and/or other repetitive sequences) by UMI-tools, the remaining reads were aligned to the human GRCh38/hg38 reference genome utilizing STAR: v. 2.6.0c. Peaks of m6A modification (cluster of reads) were firstly pinpointed by CLIPper tool (https://github.com/YeoLab/clipper) and reproducibility of clusters among biological triplicates of each sample was determined by IDR analysis (15). Log_2_ fold change was calculated in two steps: (i) log_2_ fold change of IP relative to its corresponding input for each sample, (ii) log2 fold change of cisplatin-treated samples relative to the corresponding DMSO control, and vice versa. Further analysis for determination of crosslink sites at a single nucleotide resolution in IP-enriched samples relative to their inputs was performed by PureCLIP: v. 1.3.1 (13). Validation of crosslink sites was evaluated through both reproducibility of m6A sites between three replicates and determination of DRACH motif in which m6A sites tend to be situated (16). Finally, each crosslink site was appointed to the gene and feature type based on the GENCODE Release 32 (GRCh38.p13). Further analyses of determined feature types and crosslink sites such as relative frequency of peaks that map to each feature, Metagene Plot, Metaintron Plot, and Homer Motif analysis were carried out using R tools.

Differentially methylated RNAs were further interrogated by Gene Ontology (GO) enrichment analysis, KEGG Enrichment analysis, and Reactome Pathway platform (The Gene Ontology Consortium, 2019, 17).

### SELECT

A single-base elongation and ligation-based qPCR amplification method, termed SELECT, was used to validate m^6^A miCLIP-seq data (18). Firstly, trace genomic DNA contamination was eliminated from total RNAs using Ambion™ TURBO™ DNase kit (ThermoFisher Scientific, United States) according to the manufacturer’s instructions. 1500 ng DNAse-treated RNA was annealed with 800 nM UpPrimer, 800 nM Down Primer, 100 μM dTTP in 1X CutSmart® Buffer (New England Biolabs, United States) in a total volume of 17 μl at 90 °C for 1 minute, 80 °C for 1 minute, 70 °C for 1 minute, 60 °C for 1 minute, 50 °C for 1 minute, and 40 °C for 6 minutes. This step was performed for *PHLDA1, PMAIP1, TRAP1, PIDD1* oligos and β-actin (Table 2) as control. In the second step, an enzyme mixture of 3 μl in volume that contains 0.01 U Bst 2.0 DNA polymerase (New England Biolabs, United States), 0.5 U SplintR ligase diluted with Diluent A (New England Biolabs, United States) and 10 mM adenosine triphosphate (New England Biolabs, United States) was added to the 17 μl solution from the first step. The resulting 20 μl mixture was incubated first at 40 °C for 20 minutes, then at 80 °C for 20 minutes and finally at +4 °C. qPCR analysis of the ligated SELECT products was performed by using Ampliqon RealQ Plus 2x master mix green (without ROX, Denmark) and 4 μM SELECT primer and run using the following conditions for 60 cycles: 95 °C for 5 minutes; 95 °C for 10 seconds; and 60 °C for 45 seconds. The SELECT products of the indicated site were normalized by β-actin.

**Table 2.**
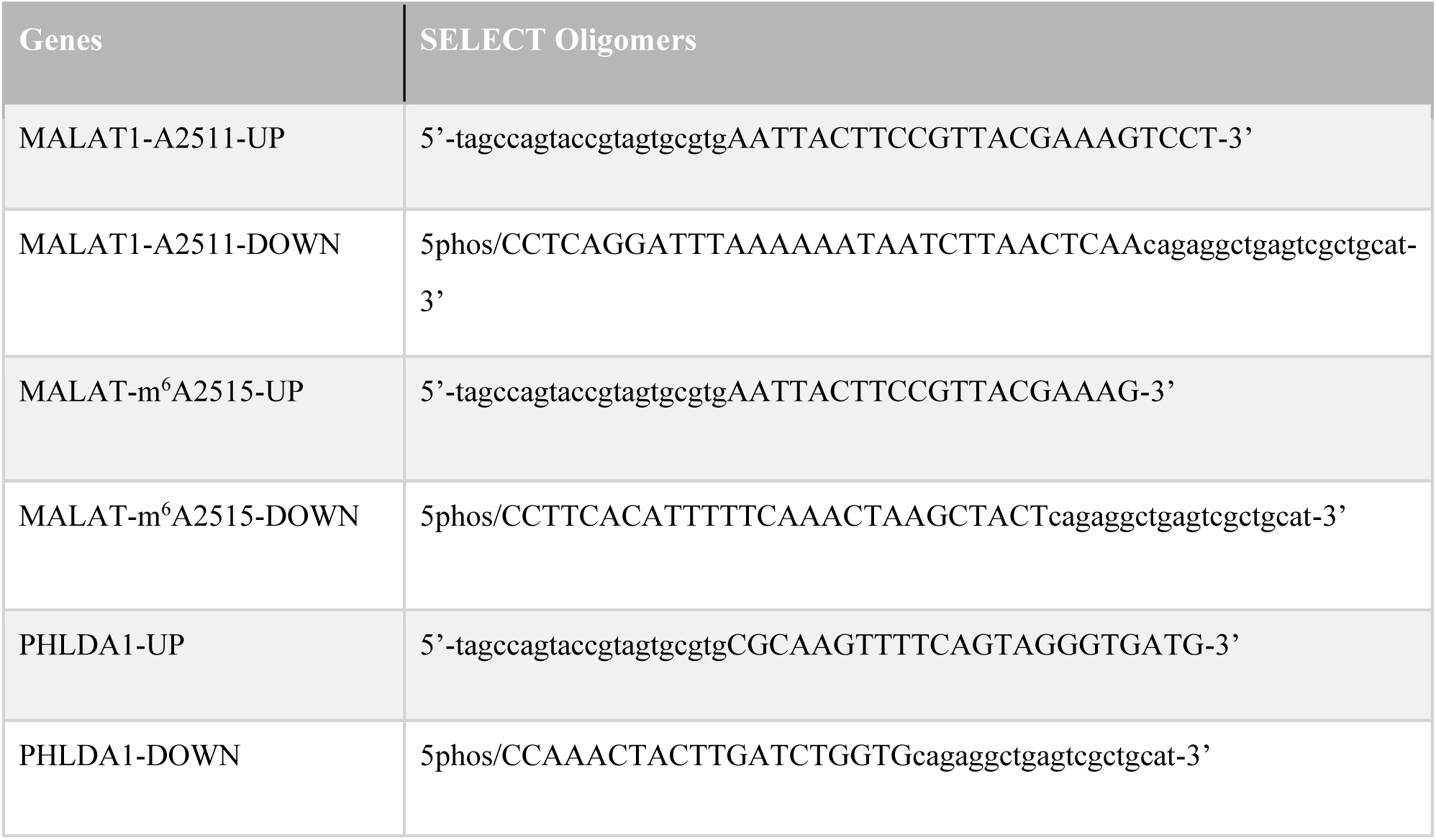
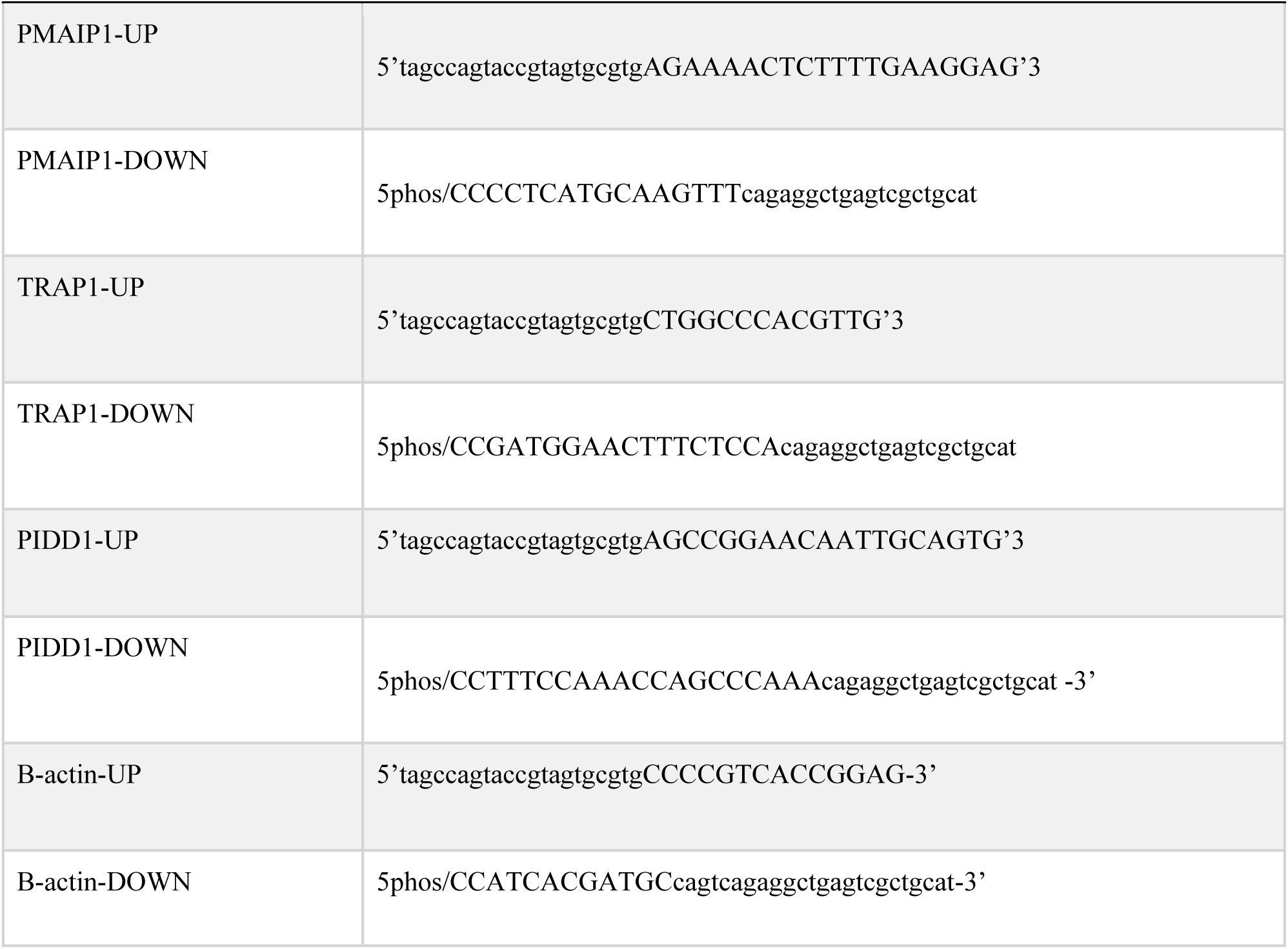
The list of oligomer sequences used in SELECT experiments

### Western Blotting

Total protein was extracted from cells using RIPA buffer (CST, United States) Subsequently, 25 μg total protein extract was fractionated on a 10% polyacrylamide gel and transferred onto PVDF membranes (ThermoFisher Scientific, United States). The membrane was blocked with 5% non-fat dry milk in 0.2% Tween-20 in Tris-buffered saline for 1h at room temperature. The primary antibodies for METTL3, METTL14, FTO and RBM15 (CST, United States) and and secondary antibodies were used in 1:1000 and 1:10000 dilution, respectively. The chemiluminescent signals were quantitated by the ImageJ program where each signal was normalized to β-actin.

### Polysome Analysis

Polysome profiling was performed according to a published procedure (19). Cells were centrifuged at 1200 RPM for 10 mins. The cells were homogenized in lysis buffer [100 mM NaCl, 10 mM MgCl_2_, 30 mM Tris-HCl (pH 7), 1% Triton X-100, 1% NaDOC, 100 μg/mL cycloheximide (Applichem, Germany) and 30 U/mL RNase Inhibitor (Promega, United States)]. The cell pellets were sheared to homogenization by passing the lysate through a 26 G needle at least 15 times and incubated on ice for 8 minutes. The lysates were then centrifuged at 12,000 *g* at 4ºC for 8 minutes. The supernatants were layered over 5-70% (w/v) sucrose gradients [100 mM NaCl, 10 mM MgCl_2_, 30 mM Tris-HCl (pH 7.0), 200 U RNase inhibitor (Promega, United States)] prepared by using an ISCO Teledyne (United States) density gradient fractionation system and centrifuged at 27,000 RPM for 2 h 55 min at 4ºC in a Beckman SW28 rotor. Fractions were collected using the ISCO Teledyne density gradient fractionation system while reading absorbance at A_254_. Fractions were pooled as mRNP, monosomal, light and heavy polysomal sub-groups based on A_254_ readings. Total RNA was phonel-extracted from fractions by using phenol-chloroform-isoamyl alcohol (25:24:1) (Applichem, Germany).

## Results

### Cisplatin regulates the expression of the m6A methylation machinery

Cisplatin is a cancer chemotherapeutic drug that is widely used as a universal inducer of apoptosis (20). We reported previously that cisplatin at a concentration of 80 μM is sufficient to trigger approximately 50% of apoptosis in HeLa cells (21). Thus, we treated HeLa cells with 80 μM cisplatin for 16 h to attain an early apoptotic rate of 55% (Figure 1A-B). We then analysed the expression levels of major genes involved in m6A RNA methylation to assess the impact of cisplatin on the m^6^A methylation machinery. To this extent, we examined the changes in the transcript levels of *METTL3, METTL14, WTAP, RBM15, FTO* and *ALKBH5* in cisplatin-treated HeLa cells. Interestingly, cisplatin treatment led to a 3.3- and 6.6-fold reduction in the mRNA levels of *METTL14* and *FTO*, respectively as determined by qPCR analyses (Figure 1C). We then checked the amount of some of these key proteins. Our western blot analyses showed downregulation of METTL14 in parallel to the qPCR analyses (Figure 1D-E). Although we did not detect any changes in the mRNA levels of *METTL3* and *RBM15* (Fig 1C), we observed a decrease in the protein levels of RBM15 and METTL3 by 2.8- and 1.4-fold, respectively (Figure 1D-E).

**Figure 1.**
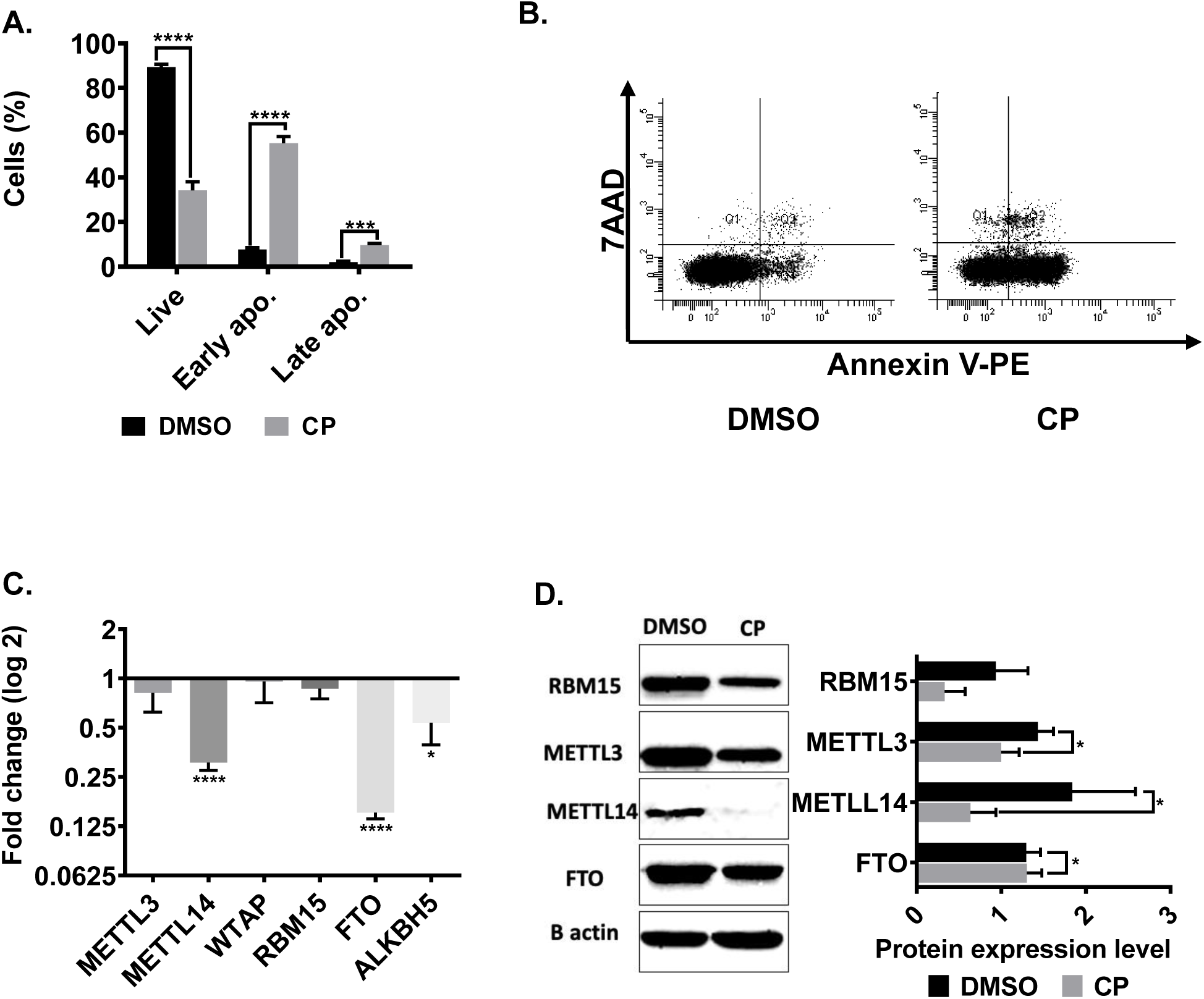
Expression patterns of m6A enzymes in cisplatin-treated HeLa cells. 1×10^6^ HeLa cells were treated with 80 μM cisplatin (CP) and 0.1% (v/v) DMSO for 16 h. The cells were stained with Annexin V/7AAD and examined by flow cytometry. Population distributions of DMSO- and CP-treated HeLa cells are depicted (**A)** Percentage of live (34%), early (55%) and late apoptosis (9.9%) and (**B)** Dot-blot analysis by flow cytometry after staining with Annexin V-PE and 7AAD. **C**. qPCR analysis of gene expression. **D**. Western blot analysis. Experiments were conducted in triplicates. *: p≤0.05, ****:p≤0.0001 by two-tailed unpaired t-test.

### Cisplatin modulates major changes in the m6A RNA methylome

Cisplatin-mediated changes in the mRNA and protein levels of key RNA methylation enzymes suggest substantial perturbations in the cellular RNA m^6^A methylome under apoptotic conditions. Therefore, we employed a genomewide approach to identify the key RNA m^6^A methylation events in cisplatin-treated HeLa cells. To this extent, we performed an m^6^A miCLIP-seq analysis with three biological replicates of total RNAs isolated from DMSO- and cisplatin-treated HeLa cells. We normalized the changes in RNA m^6^A methylation events against transcript abundance to obtain the differentially m^6^A-methylated transcripts. Our analyses revealed changes in the m^6^A methylation pattern at 7658 adenosine residues with an average number of 12 m^6^A methylation per transcript (Supplementary Table 1). The extent of m^6^A methylation was reduced at 6236 sites while we detected induction in the m^6^A RNA methylation pattern of 1422 sites. We then generated metaintron plots to interrogate the distribution of m^6^A sites throughout transcripts. We noticed an enrichment at the 5’ and 3’ UTRs of transcripts with increased and decreased m^6^A residues, respectively (Figure 2A-B). Additionally, down-regulated m^6^A methylation was predominantly localized to the 3’ UTR and coding sequence of mRNAs, and the up-regulated m6A residues have homogenic distribution throughout mRNA (Figure 2C-D). To uncover which transcripts are specifically targeted by the m^6^A methylation machinery under cisplatin-induced apoptotic conditions, we carried out gene ontology analyses with differentially m^6^A-methylated transcripts (Supplementary Figure 2). Strikingly, as many as 132 genes associated with apoptosis were subject to differential m^6^A methylation (Figure 2E, Supplementary Table 3). An IGV screen shot of one of the methylation sites is presented in Figure 2F.

**Figure 2.**
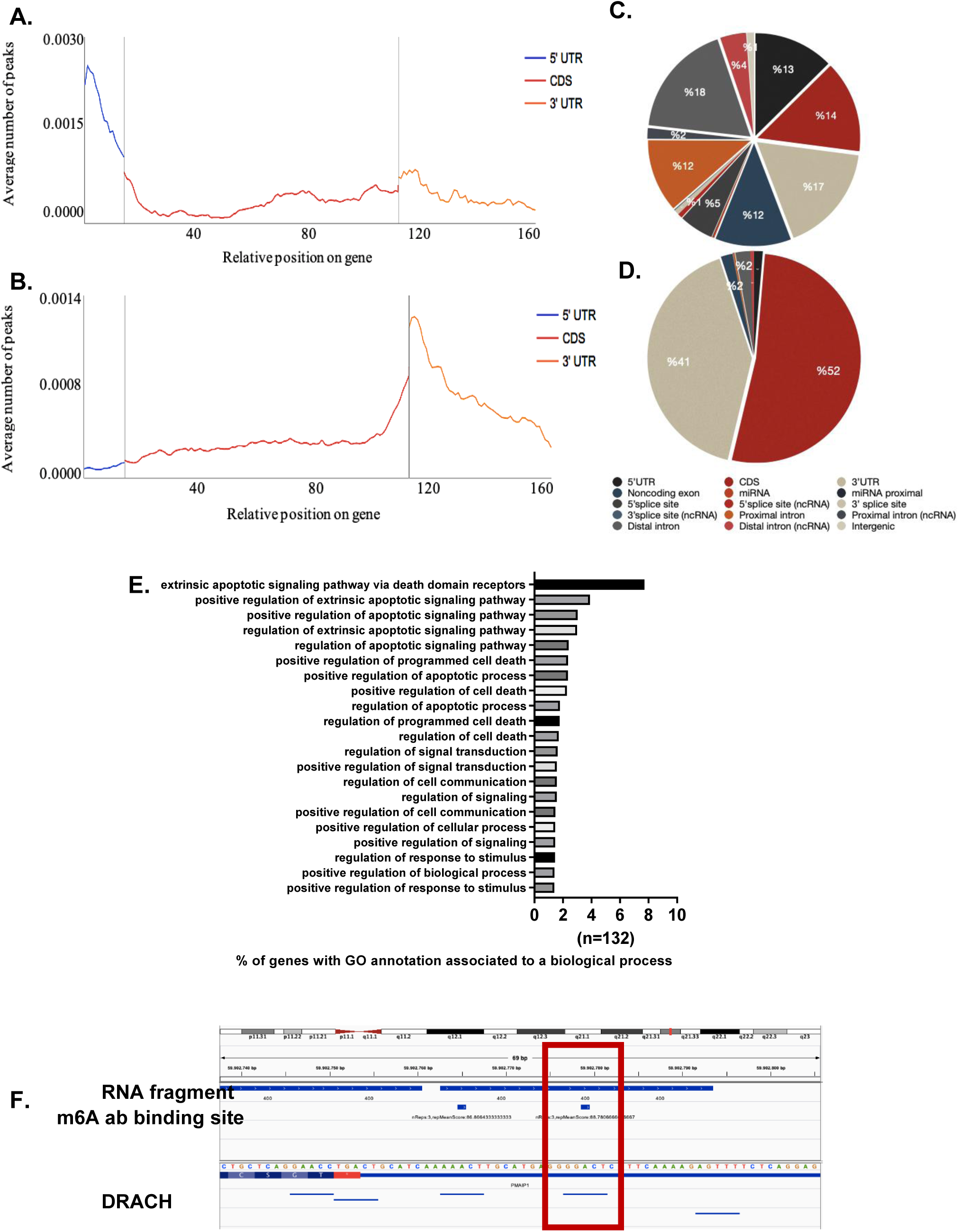
m^6^A RNA methylome profile of cisplatin-treated HeLa cells. The m6A methylome of DMSO control and CP-treated HeLa cells was obtained by miCLIP-seq as described in Materials and Methods. Distribution of upregulated (**A)** and downregulated **(B)** m^6^A peaks are shown across all transcripts. Pie chart of upregulated (**C)** and downregulated **(D)** m^6^A peaks representing their location on transcripts. Biological replicates: n = 3 per group. **E**. Gene ontology (GO) analysis of differentially m^6^A-methylated transcripts associated with apoptosis. All biological processes were plotted having a false discovery rate (FDR) < 0.05. **F**. 524^th^ adenosine residue in PMAIP1.

Although miCLIP-seq is a highly sensitive approach to examine m^6^A-methylated transcripts genomewide, the potential for false positive sites that might arise from a cross-hybridization mandates validation of m^6^A sites of interest (16). Thus, we employed SELECT (18) to confirm differential m^6^A methylation on four mRNAs, namely *PHLDA1, PIDD1, PMAIP1* and *TRAP1*, all of which have been reported to be key players in modulation of apoptosis [Table 1; (4,22–24)]. To this extent, we first used the m^6^A site at the 2525^th^ adenosine nucleotide of *MALAT1* as a positive control to ensure that this method properly detects m^6^A-methylated residues (18). Our results showed a 1.85 Cq difference in the amplification of the ligated product in METTL3-knockdown HeLa cells (Figure 3A-B), suggesting the proper detection of m^6^A residues. We detected no difference in the amplification of the ligated product flanking the 2511^th^ adenosine residue, which is known not to harbour any m^6^A, further confirming the validity of SELECT (Figure 3B). We then used the same approach to check the presence of an m6A residue on *PHLDA1, PIDD1, PMAIP1* and *TRAP1* mRNAs. Based on the amount of the amplified ligation product, we verified the m6A residues on *PMAIP1* and *TRAP1* mRNAs (Figure 3C). A similar amplification efficiency of GAPDH without m^6^A was used as a negative control in the same experimental setting to demonstrate the dependency of the amplification product on METTL3-mediated m6A methylation (Figure 3C).

**Figure 3.**
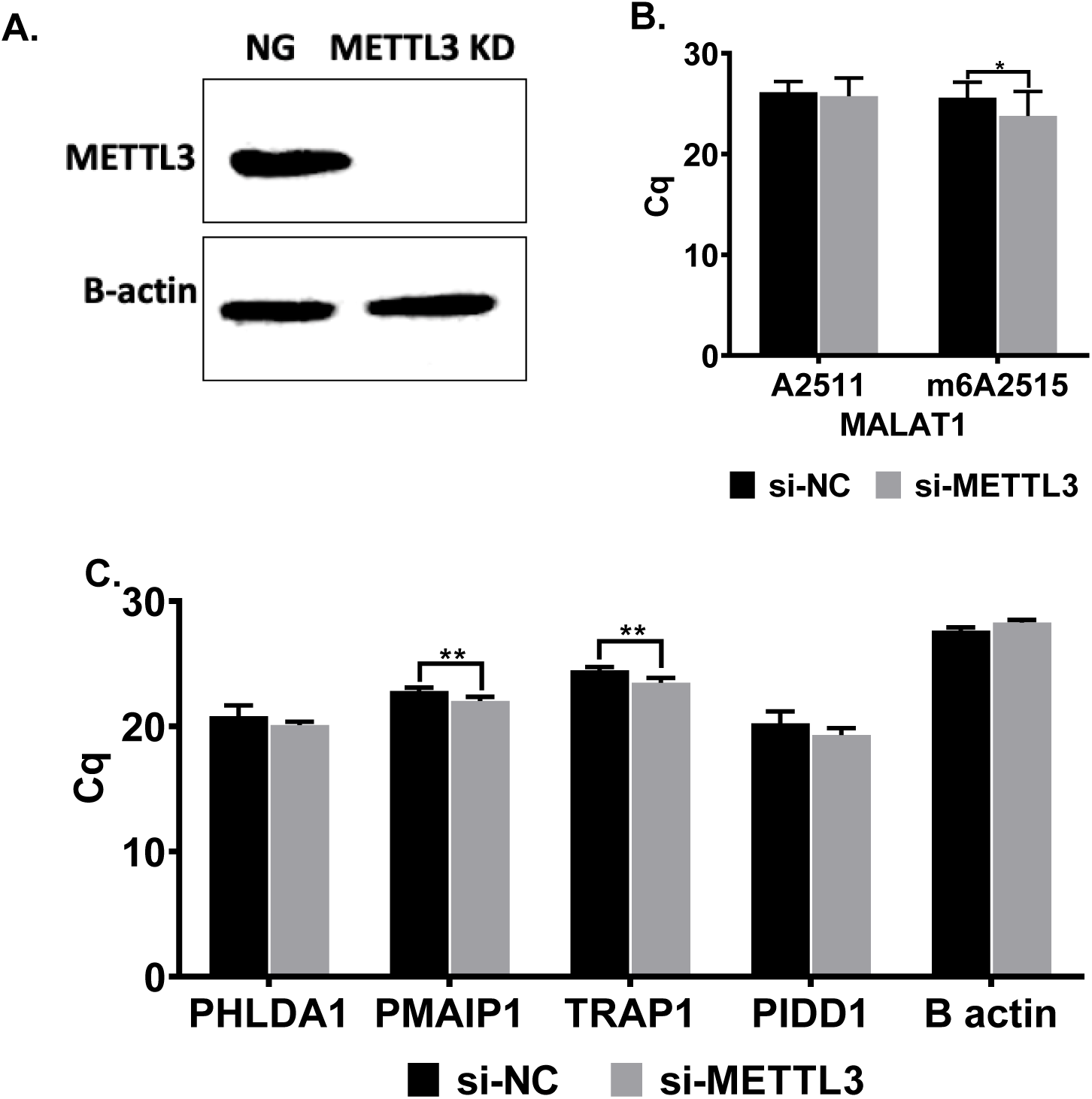
Validation of m^6^A residues in cisplatin-treated HeLa cells. **A.** 0.6×10^6^ HeLa cells were transfected with 25 nM negative (NG) or METTL3 siRNA (siMETTL3) in 10-cm culture dishes for 72 h. Western blot was performed to examine the efficiency of knockdown. Beta actin was used as loading control. **B**. SELECT qPCR Rn/ CT values of 2515^th^ m^6^A site in MALAT1 as a positive control and 2511^th^ A in MALAT1 as a negative control. (n=3, two-tailed unpaired t-test). **C**. SELECT qPCR Rn/CT values of the 904^th^ m^6^A in PHLDA1, 524^th^ m^6^A in PMAIP1 sites, 1370^th^ m^6^A in PIDD1 and 44419^th^ m^6^A in TRAP1 in METTL3 KD HeLa cells (n=3, two-tailed unpaired t-test). Experiments were performed in triplicates. p>0.05, *: p≤0.05, **: p≤0.01.

Verification of m^6^A sites on candidate mRNAs has prompted us to examine the extent of m^6^A methylation under cisplatin-induced apoptotic conditions in HeLa cells. To this end, we treated HeLa cells with cisplatin for 16 h and conducted SELECT experiments. Our analyses revealed cisplatin-mediated increases in the m6A methylation of *PHLDA1, PMAIP1* and *TRAP1* mRNAs by a Cq difference of 2.4, 1.1 and 1.6, respectively (Figure 4A-C). As apoptosis is a highly dynamic process (1), we also interrogated the time kinetics of the m6A methylation status of these candidates. We observed a transcript-specific methylation kinetics. For example, for *PHLDA1* and *PMAIP1*, we detected a sudden increase in methylation at 8 h followed by relative less increase in 16 and 32 h (Figure 4A-B). *TRAP1* experienced a decrease in the methylation rate at 8 h followed by an elevation at 16 and 32 h (Figure 4C). We were unable to validate any cisplatin-induced m^6^A changes on *PIDD1* (Figure 4D). (1370^th^ adenosine in PHLDA1, 524^th^ adenosine in PMAIP1, 904^th^ adenosine in PIDD1 and 44419^th^ adenosine in TRAP1). Subsequently, we examined the effect of cisplatin on the same m^6^A residues in ME-180 and MCF7 cells to inspect the cell specificity. Under the same cisplatin treatment conditions, we observed similar m^6^A pattern with CP treated HeLa cells. (Figure 5A-B).

**Figure 4.**
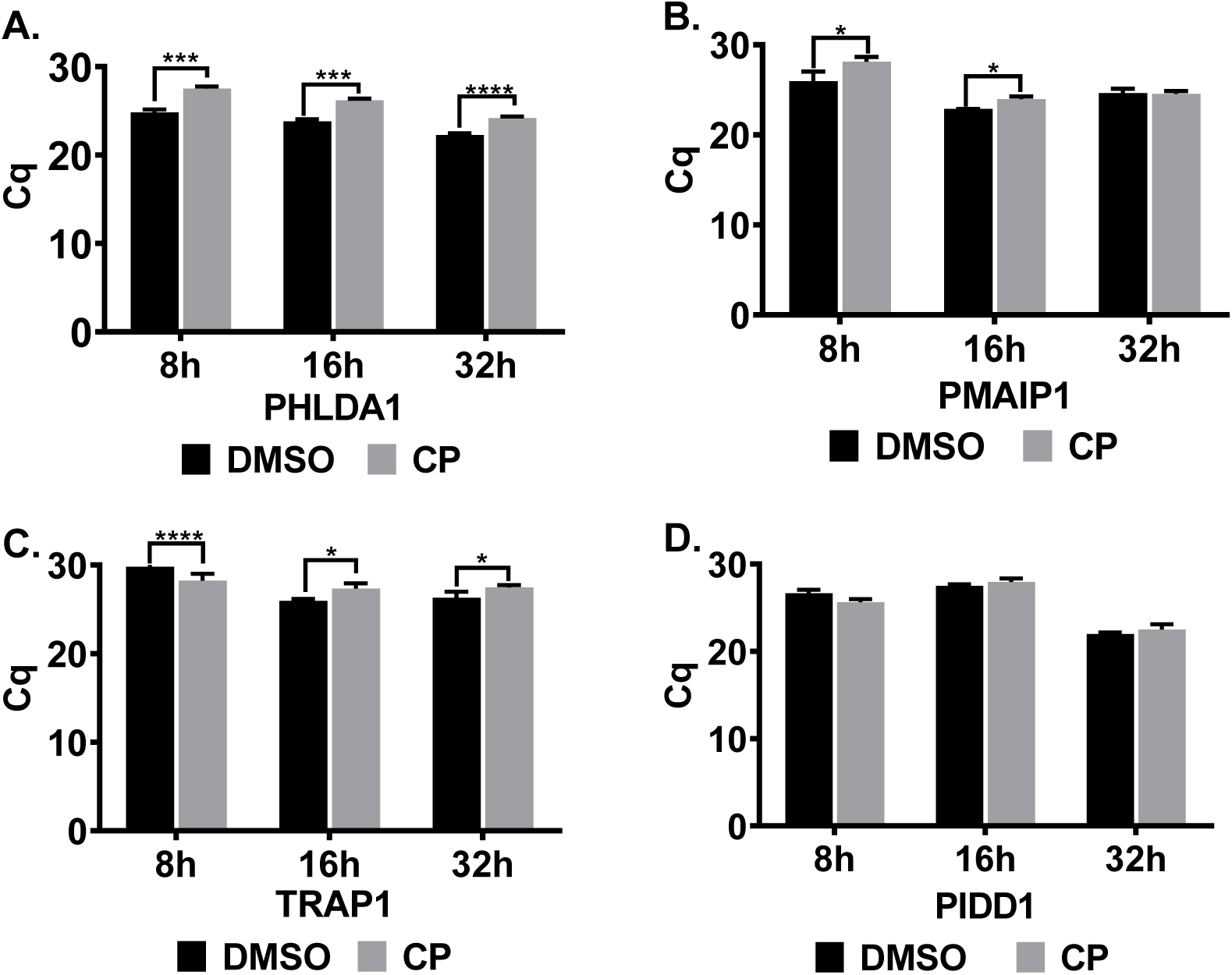
Dynamic m^6^A methylation in cisplatin-treated HeLa cells. 1×10^6^ HeLa cells were treated with 80 μM CP- and 0.1% (v/v) DMSO for 8h, 16 h and 32h. SELECT qPCR were used to quantitate the Rn/CT values of the 904^th^ m6A in PHLDA1, 524^th^ m6A in PMAIP1, 1370^th^ m6A in PIDD1 and 44419^th^ m6A in TRAP1 using total RNAs isolated from DMSO- and CP-treated cells. Error bars represent SD for 3 biological x 2 technical replicates p>0.05, *: p≤0.05, ***: p≤0.001, ****: p≤0.0001 by two-tailed unpaired t-test.

**Figure 5.**
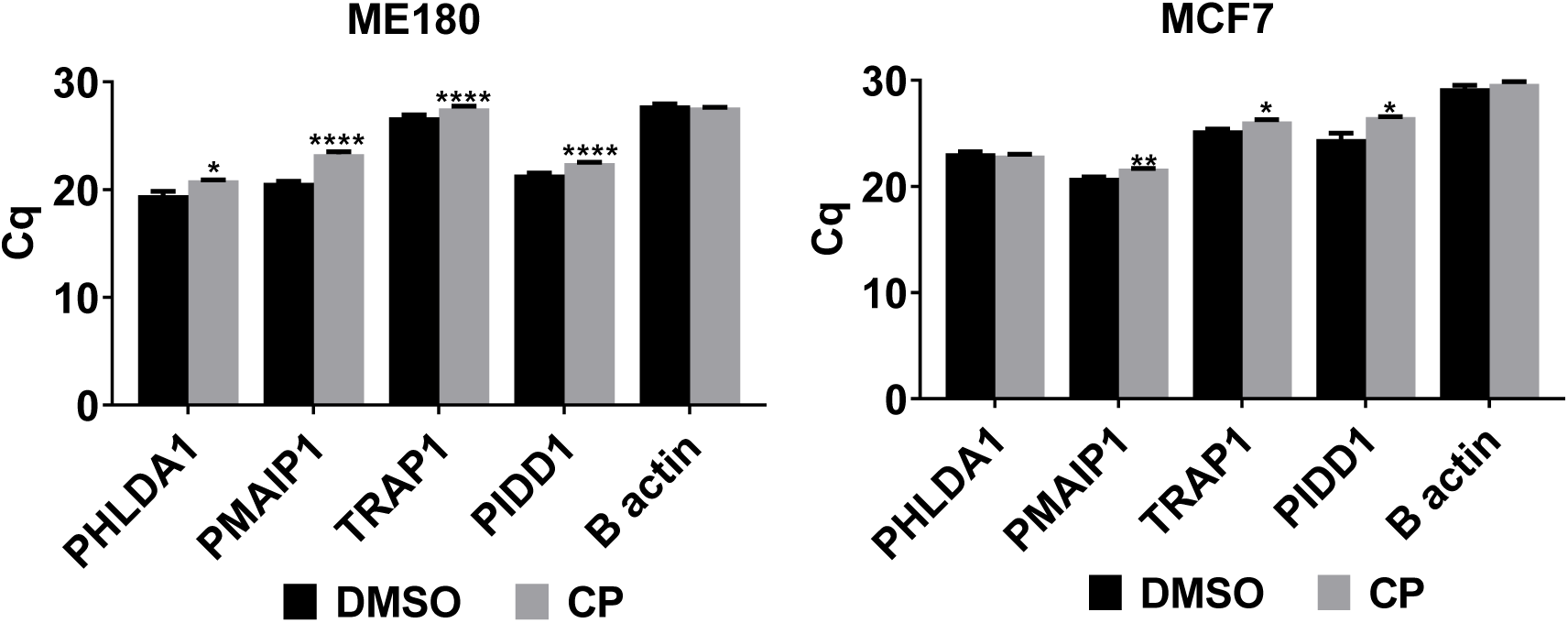
m^6^A methylation in apoptotic ME180 and MCF7 cells. 1.2×10^6^ ME180 and MCF7 cells were treated 80 μM and 50 μM CP, respectively, and 0.1% (v/v) DMSO as control for 16 h. SELECT qPCR were run to obtain the Rn/CT values of m6A residues in PHLDA1, PMAIP1, PIDD1 and TRAP1 in CP treated ME180 and MCF7 cells. Beta actin was used for normalization. ns: non-significant, p>0.05, *: p≤0.05, **: p≤0.01, ****: p≤0.0001 by two-tailed unpaired t-test.

### Cisplatin modulates translational efficiency of mRNAs in a METTL3-dependent manner

RNA m6A methylation may regulate the fate of transcripts both transcriptionally and posttranscriptionally (25). Thus, we first examined the impact of RNA m^6^A methylation on transcript abundance in the presence or absence of cisplatin and/or METTL3. Although *METTL3* knockdown did not change the viability of HeLa cells in the absence of cisplatin (e.g., DMSO treatment), we observed a 3.5 % decrease in the percentage of live cells and a 5.8% increase in the percentage of early apoptotic cells (Figure 1A, *P* < 0.05), suggesting that *METTL3* knockdown probably sensitizes HeLa cells to cisplatin-induced apoptosis. Before we investigated the cisplatin inducibility of candidate genes in the absence of METTL3, we first interrogated their abundance in cisplatin-treated and *METTL3* knockdown HeLa cells separately. Based on our qPCR analyses, cisplatin treatment led to a 14.3-12.1- and 3.2-fold increase in the transcript amount of *PHLDA1, PMAIP1* and *PIDD1*, respectively (Figure 6B). On the other hand, *METTL3* knockdown did not have a major impact on the transcript abundance of any of the candidates tested (Figure 6C). We then measured the mRNA amounts in DMSO- and cisplatin-treated HeLa cells following the knockdown of METTL3. Similarly, no major effects were observed on transcript abundance (Figure 6C-D).

**Figure 6.**
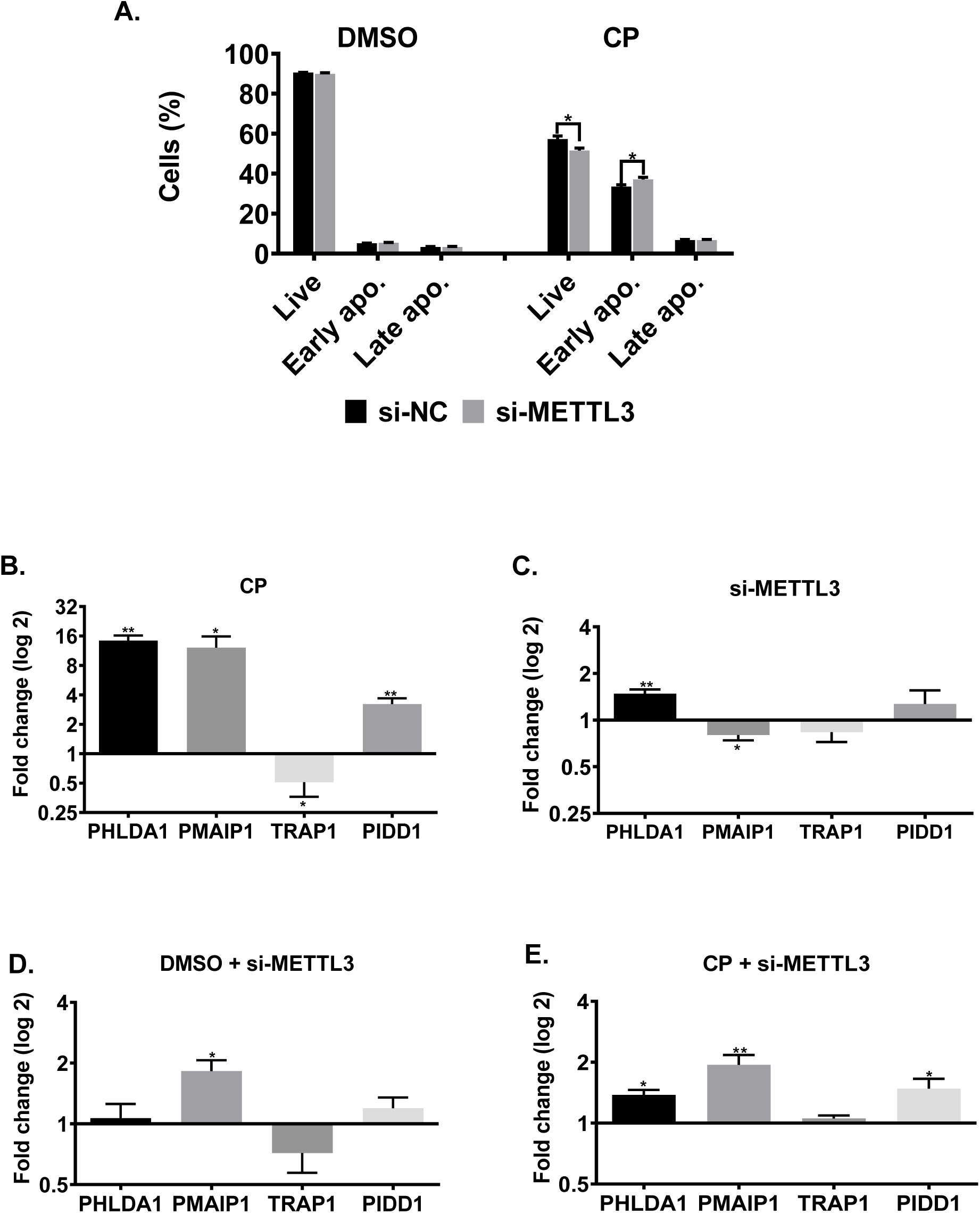
RNA abundance in cisplatin-treated METTL3 knockdown HeLa cells. **A.** HeLa cells were treated with 40 μM CP for 16 h following siRNA transfection. DMSO (0.05%) was used as negative control. qPCR analyses of total RNAs isolated from DMSO and CP-treated **(B)**, si-NG and si-METTL3-transfedted cells **(C)**, si-METTL3-transfected and DMSO-treated cells **(D)** and si-METTL3-transfected and CP-traeted cells **(E)**. p>0.05, *: p≤0.05, **: p≤0.01 by two-tailed unpaired t-test.

As the transcript abundance of candidate genes does not appear to be influenced by METTL3 knockdown in cisplatin-treated HeLa cells, we hypothesized that RNA m^6^A methylation could be critical for translational efficiency of target RNAs under cisplatin-induced apoptotic conditions. Thus, we examined the polysome profiles of cells under various conditions as the association of mRNAs with polysomes is a good indicator of their translational efficiency (26). Knockdown of *METTL3* did not appear to have a discernible change in the overall polysome profile (Figure 7A), excluding any global effect on translation. We then investigated the association of individual mRNAs with actively translating polysomes. To this extent, we first fractionated the cytoplasmic extracts of HeLa cells into four major fractions (1) mRNP fraction composed primarily of non-translated ribonucleoproteins; (2) monosome constituting mRNAs associated with a single ribosome; (3) light polysome that contains mRNAs associated with up to 3 ribosomes; and (4) heavy polysome that contains mRNAs with more than 3 ribosomes. We performed qPCR analyses with total RNAs phenol extracted from each fraction. Our results exhibited almost no changes in the translational efficiency of *PHLDA1, PIDD1, PMAIP1* and *TRAP1* mRNAs upon *METTL3* knockdown (Figure 7B-E). We then examined the translational efficiency of these transcripts in cisplatin-treated HeLa cells upon *METTL3* knockdown to interrogate the potential contribution of METTL3 to the translational efficiency of transcripts under cisplatin-induced apoptosis. In agreement with our data in Figure 7, the treatment of HeLa cells with control DMSO did not cause any changes in the sedimentation of the transcripts on sucrose gradients upon METTL3 knockdown (Figure 8 A,C,E and G). Interestingly, we detected CP-mediated changes in the translational efficiency of *PHLDA1* (Figure 8B, monosome, 7.1-fold, *P*< 0.05), *PMAIP1* (Figure 8D, light polysome, 14.2-fold, *P*< 0.05) and *PIDD1* (Figure 8H, mRNP, 2.4-fold, *P*< 0.05) mRNAs. CP-mediated increase in the translational efficiency of *PMAIP1* has stricken our attention as PMAIP1 has been reported to mediate apoptosis by inducing the intrinsic apoptotic pathway (27). To substantiate our observation with the polysome analyses, we measured the protein amount of PMAIP1 in CP-treated HeLa cells upon *METTL3* knockdown. PMAIP1 was undetectable in DMSO-treated cells. However, CP caused a sharp increase in the protein level (Figure 9A). We then examined the protein level of PMAIP1 in *METTL3* knockdown cells upon CP treatment. Interestingly, *METTL3* knockdown resulted in an increase in the protein level of PMAIP1 compared to the cells transfected with the control siRNA (Figure 9A-B, 1.5-fold, P<0,001). The CP-mediated increase in the protein level of PMAIP1 in *METTL3-*knockdown cells was accompanied with a parallel increase in the amount of cleaved caspase 9 (Figure 9C). Since *PMAIP1* is a p53-responsive gene (27), we also analysed the effect of *METTL3* knockdown on the protein level of p53. Strikingly, METTL3 knockdown resulted in a dramatic reduction in the abundance of P53 (Figure 9C).

**Figure 7.**
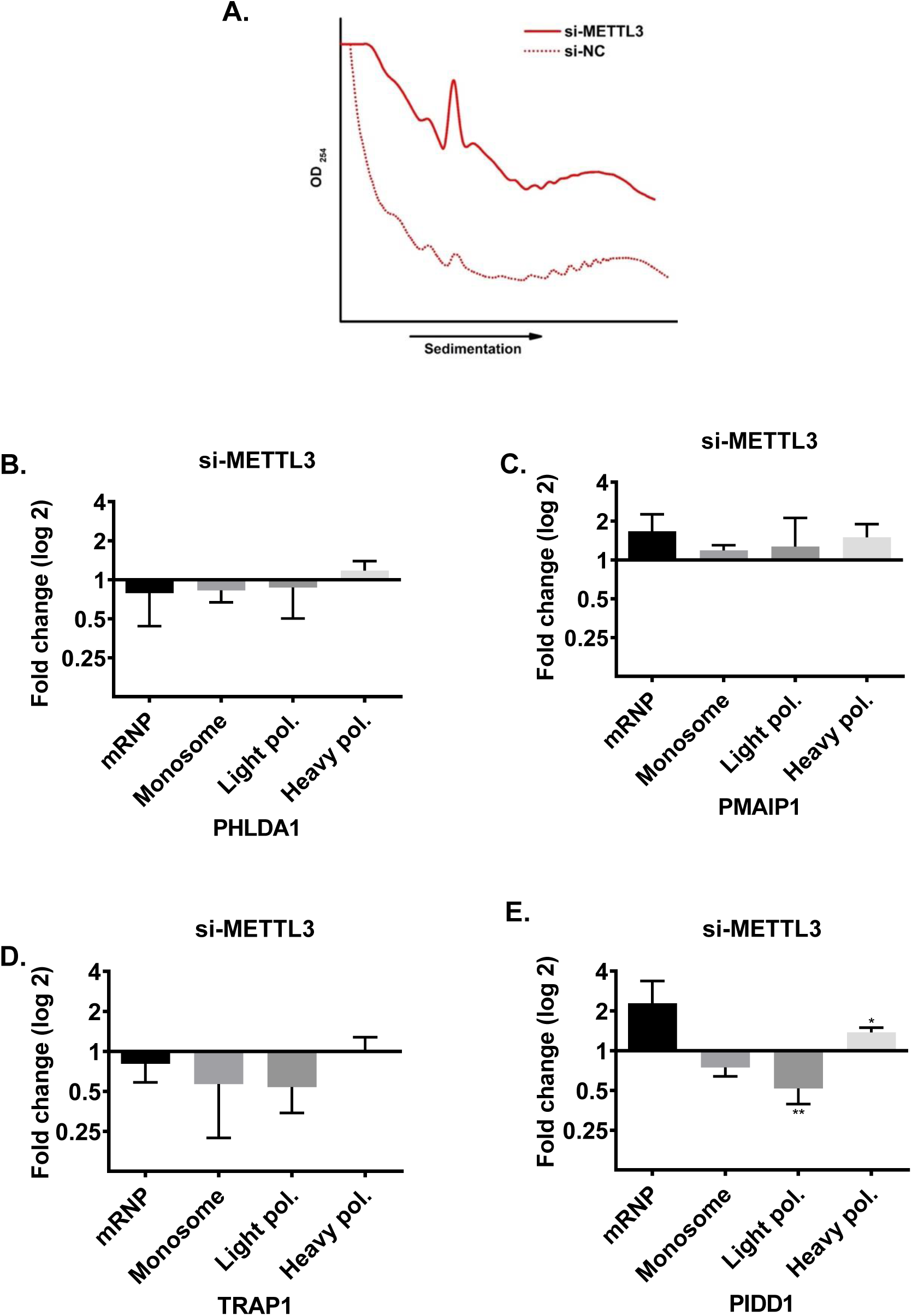
Polysome profiling in METTL3 knockdown HeLa cells. **A.** Polysome profiles of cells transfected with si-METTL3 or negative siRNA (si-NC). Total RNAs were phenol-extracted from each fraction collected based on the polysome profile and transcript abundance was measured by qPCR (B-E). n=3. p>0.05, *: p≤0.05, **: p≤0.01 by two-tailed unpaired t-test.

**Figure 8.**
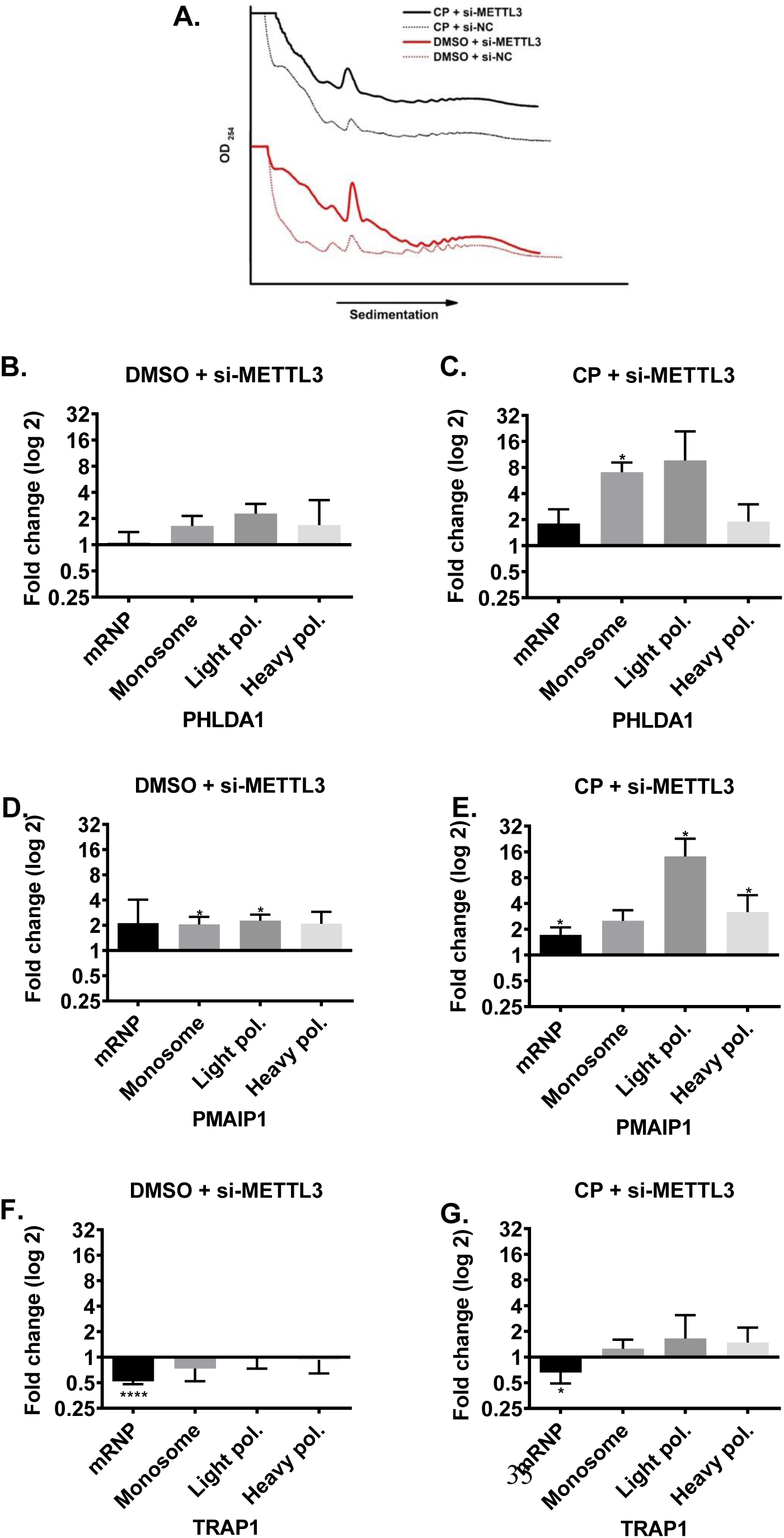
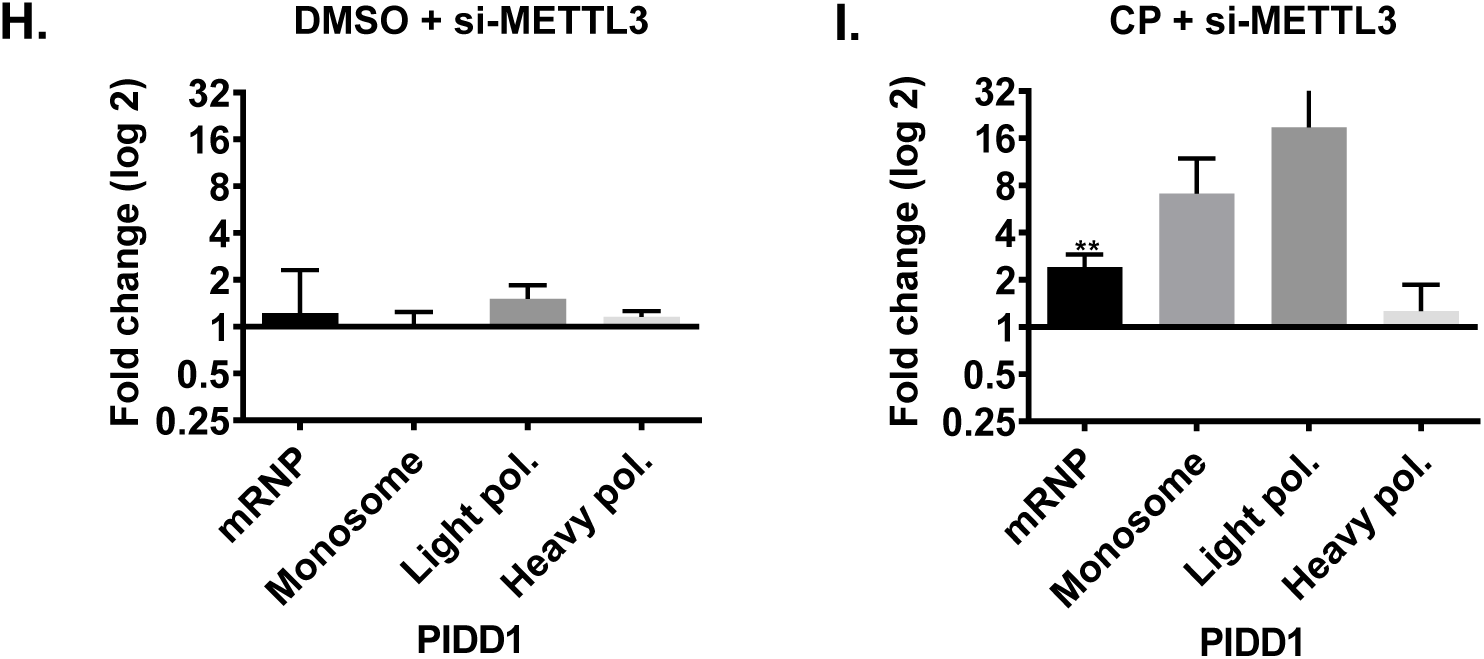
Polysome profiling in cisplatin-treated METTL3 knockdown HeLa cells. Polysome profiles of cells transfected with si-METTL3 or negative siRNA (si-NC) and treated with 40μM CP for 16h. Total RNAs were phenol-extracted from each fraction and transcript abundance was examined by qPCR (B-I). n=3. *: p≤0.05, **: p≤0.01, ****: p≤0.0001 by two-tailed unpaired t-test.

**Figure 9.**
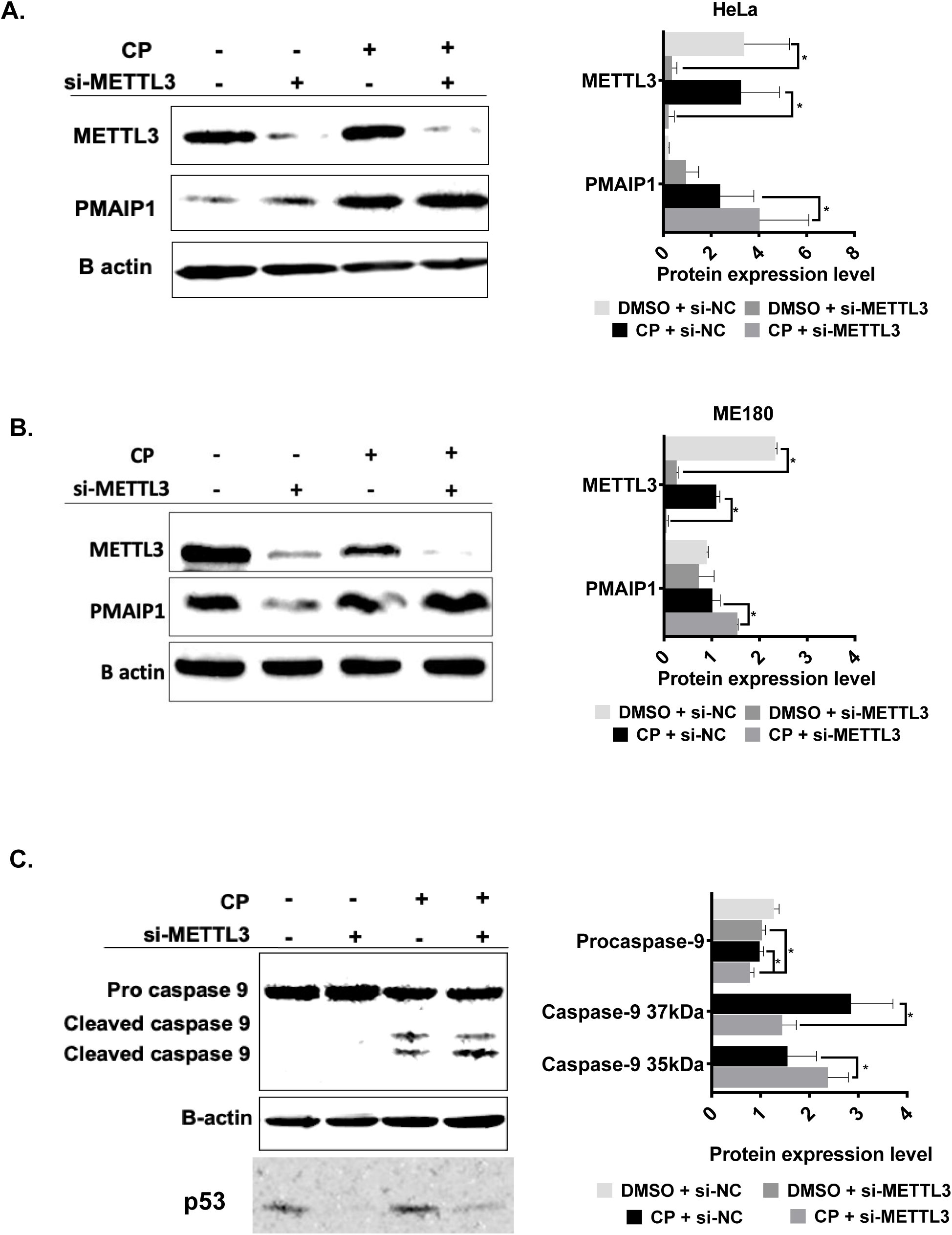
The METTL3-P53-PMAIP1 axis in METTL3 knockdown HeLa cells. Western blot analysis of total protein (25 μg) from HeLa **(A)**, ME180 **(B)** and MCF **(C)** cells transfected with si-METTL3 and treated with CP as in Figure 8. B-actin was used as loading control. n=3 *: p≤0.05 by two-tailed unpaired t-test.

## Discussion

We report for the first time that the m^6^A methylation machinery plays a fundamental role in modulating cisplatin-mediated apoptosis in HeLa cells (Figure 1). As a drug with pleiotropic effects, cisplatin induces dynamic changes in the m6A methylome associated not only with apoptosis but also with stress, growth, migration etc. (Figure 2). Our genomewide approach uncovers that the METTL3-p53-PMAIP1 axis appears to modulate apoptosis through the METTL3-mediated changes in p53 and PMAIP1 protein amounts (Figure 9).

METTL3 has been reported to play a role in cell death as its knockdown induces apoptosis in HepG2 cells by modulating the *P53* signalling and splicing of isoforms of *MDM2, FAS* and *BAX* (6). METTL3 was also shown to be involved in the selective recruitment of DNA polymerase to damaged DNA sites to orchestrate repair (28). Additionally, there are reports on the role of ALKBH5-mediated methylation on cisplatin resistance (29,30); Although we did not detect any change in the rate of apoptosis upon in *METTL3* knockdown cells, the knockdown sensitized the cells to cisplatin-induced apoptosis (Figure 3A). Our gene expression analyses in apoptotic HeLa cells also revealed major perturbations in the amounts of METTL3, METTL14 and RBM15 without any change in FTO (Figure 1). Although these observations clearly suggest a critical role for m^6^A methylation in coordinating different types of cell death (31), a complete m^6^A methylome profiling would be needed to gain insight into the extent of dynamic changes in the m^6^A RNA methylome under cisplatin-induced apoptotic conditions. We detected as many as 972 differentially m^6^A-methylated mRNAs, of which 132 were associated with apoptosis (Figure 2). Condition-specific enrichment of m^6^A on mRNAs has been reported previously. For example, m^6^A residues in 5’ UTRs have been associated with cap-independent translation (32). On the other hand, m^6^A residues on coding regions (CDs) were reported to induce translation by helping resolve secondary structures (33). We did not detect any enrichment on any specific regions of mRNAs except for a slight enrichment on the terminal part of 5’ and 3’ UTRs (Figure 2A). SELECT-based validation confirmed the m^6^A results (Figure 3C). Interestingly, cisplatin-mediated changes in m^6^A methylomes were highly dynamic. For example, m^6^A residues, which appeared on *PMAIP1* mRNA as early as 8h upon cisplatin treatment, dissipated by 32h (Figure 4B). For all genes tested, we observed a similar m^6^A methylation pattern in HeLa, ME180 and MCF7 cells (Figure 4-5).

m^6^A residues determine the fate of mRNAs at both transcriptional and posttranscriptional levels (25). We first examined the abundance of our candidate mRNAs to probe into the impact of differential m^6^A methylation on the transcription rate and/or mRNA stability. Our qPCR analyses revealed no changes in the mRNA abundance of *PHLDA1, PMAIP1, TRAP1* and *PIDD1* (Şekil 6). However, we observed a strong association between METTL3 and translational efficiency of these mRNAs especially under cisplatin treatment conditions (Figure 8). Although METTL3 is reported to promote translation in human cancer cells independent of its catalytic activity and m^6^A readers (34), *METTL3* kockdown resulted in a better association of our candidate mRNAs with polysomes, especially PHLDA1, PIDD1 and PMAIP1, under cisplatin treatment conditions (Figure 8). It is interesting that METTL3 knockdown did not influence the extent of polysome association of these RNAs under control DMSO treatment (Figure 8).

PMAIP1 is a proapoptotic protein that targets MCL1 or BCL2A1 proteins for degradation (35). As a p53-responsive gene, PMAIP1 induces apoptosis in HeLa cells by activating caspase-9 (27). Thus, we examined whether the enhanced translational efficiency of *PMAIP1* in METTL3 knockdown cells results in an increase in its protein amount. Expectedly, METTL3 knockdown leads to an increase in PMAIP1 amount both in HeLa and ME180 cells (Figure 9A-B). Additionally, we detected an elevation in the amount of cleaved caspase-9 in cisplatin-treated HeLa cells upon *METTL3* knockdown. Since *PMAIP1* is a p53-responsive gene, we interrogated whether p53 is targeted by the m^6^A methylation machinery as well. Interestingly, METTL3 knockdown resulted in a great reduction in the total P53 abundance.. Recently, it was reported that the Mettl3 epitranscriptomic writer amplifies P53 stress responses in mice (36). Mettl3 directs METTL3 to p53 downstream genes for differential methylation, including PMAIP1. It is highly interesting that PMAIP1 is targeted by the m^6^A machinery both transcriptionally (36) and posttranscriptionally (Figure 9).

## Acknowledgements

The authors would like to thank Prof. Dr. Dilek Telci Temeltaş for PMAIP1 antibody. The authors would also like to thank Özgür Okur and Murat Delman for flow cytometry analyses, and BIOMER (IZTECH, Turkey) for the instrumental help. This study was funded by the Scientific and Technological Research Council of Turkey (TÜBİTAK) (Project No: 217Z234 to BA).

## Author Contributions

BA contemplated the project. ÖT performed polysome experiments. AA, BS, ABG and İEV performed all other experiments. AA and BA wrote the manuscript, ÖT, BS, ABG and İEV read and approved the manuscript.

## References

1. Elmore S. Apoptosis: A Review of Programmed Cell Death. Toxicol Pathol. 2007;35(4):495–516.

2. Chen P, Li J, Chen YC, Qian H, Chen YJ, Su JY, et al. The functional status of DNA repair pathways determines the sensitization effect to cisplatin in non-small cell lung cancer cells. Cell Oncol. 2016;39(6):511–22.

3. Ming X, Groehler A, Michaelson-Richie ED, Villalta PW, Campbell C, Tretyakova NY. Mass Spectrometry Based Proteomics Study of Cisplatin-Induced DNA-Protein Cross-Linking in Human Fibrosarcoma (HT1080) Cells. Chem Res Toxicol. 2017;30(4):980–95.

4. Wang J, Thomas HR, Li Z, Yeo NC, Scott HE, Dang N, et al. Puma, noxa, p53, and p63 differentially mediate stress pathway induced apoptosis. Cell Death Dis [Internet]. 2021;12(7). Available from: http://dx.doi.org/10.1038/s41419-021-03902-6

5. Wei CM, Gershowitz A, Moss B. Methylated nucleotides block 5′ terminus of HeLa cell messenger RNA. Cell. 1975;4(4):379–86.

6. Dominissini D, Moshitch-Moshkovitz S, Schwartz S, Salmon-Divon M, Ungar L, Osenberg S, et al. Topology of the human and mouse m6A RNA methylomes revealed by m6A-seq. Nature. 2012;485(7397):201–6.

7. Meyer KD, Saletore Y, Zumbo P, Elemento O, Mason CE, Jaffrey SR. Comprehensive analysis of mRNA methylation reveals enrichment in 3′ UTRs and near stop codons. Cell. 2012;149(7):1635–46.

8. Vu LP, Pickering BF, Cheng Y, Zaccara S, Nguyen D, Minuesa G, et al. The N 6 - methyladenosine (m 6 A)-forming enzyme METTL3 controls myeloid differentiation of normal hematopoietic and leukemia cells. Nat Med. 2017;23(11):1369–76.

9. Wang H, Xu B, Shi J. N6-methyladenosine METTL3 promotes the breast cancer progression via targeting Bcl-2. Gene [Internet]. 2020;722(June 2019):144076. Available from: https://doi.org/10.1016/j.gene.2019.144076

10. Huang Y, Su R, Sheng Y, Dong L, Dong Z, Xu H, et al. Small-Molecule Targeting of Oncogenic FTO Demethylase in Acute Myeloid Leukemia. Cancer Cell [Internet]. 2019;35(4):677-691.e10. Available from: https://doi.org/10.1016/j.ccell.2019.03.006

11. Xu F, Li CH, Wong CH, Chen GG, Lai PBS, Shao S, et al. Genome-wide screening and functional analysis identifies tumor suppressor long noncoding RNAs epigenetically silenced in hepatocellular carcinoma. Cancer Res. 2019;79(7):1305–17.

12. Yaylak B, Erdogan I, Akgul B. Transcriptomics analysis of circular RNAs differentially expressed in apoptotic HeLa cells. Front Genet. 2019;10(MAR):1–10.

13. Krakau S, Richard H, Marsico A. PureCLIP: Capturing target-specific protein-RNA interaction footprints from single-nucleotide CLIP-seq data. Genome Biol. 2017;18(1):1–17.

14. Smith T, Heger A, Sudbery I. UMI-tools: Modeling sequencing errors in Unique Molecular Identifiers to improve quantification accuracy. Genome Res. 2017;27(3):491–9.

15. Li Q, Brown JB, Huang H, Bickel PJ. Measuring reproducibility of high-throughput experiments. Ann Appl Stat. 2011;5(3):1752–79.

16. Linder B, Grozhik A V., Olarerin-George AO, Meydan C, Mason CE, Jaffrey SR. Single-nucleotide-resolution mapping of m6A and m6Am throughout the transcriptome. Nat Methods. 2015;12(8):767–72.

17. Griss J, Viteri G, Sidiropoulos K, Nguyen V, Fabregat A, Hermjakob H. ReactomeGSA - Efficient Multi-Omics Comparative Pathway Analysis. Mol Cell Proteomics. 2020;19(12):2115–24.

18. Xiao Y, Wang Y, Tang Q, Wei L, Zhang X, Jia G. An Elongation- and Ligation-Based qPCR Amplification Method for the Radiolabeling-Free Detection of Locus-Specific N6-Methyladenosine Modification. Angew Chemie - Int Ed. 2018;57(49):15995–6000.

19. Göktaş Ç, Yiğit H, Coşacak Mİ, Akgül B. Differentially expressed tRNA-derived small RNAs co-sediment primarily with non-polysomal fractions in Drosophila. Genes (Basel). 2017;8(11).

20. Kelland L. The resurgence of platinum-based cancer chemotherapy. Nat Rev Cancer. 2007;7(8):573–84.

21. Gurer DC, Erdogan İ, Ahmadov U, Basol M, Sweef O, Cakan-Akdogan G, et al. Transcriptomics Profiling Identifies Cisplatin-Inducible Death Receptor 5 Antisense Long Non-coding RNA as a Modulator of Proliferation and Metastasis in HeLa Cells. Front Cell Dev Biol. 2021;9(August):1–13.

22. Murata T, Sato T, Kamoda T, Moriyama H, Kumazawa Y, Hanada N. Differential susceptibility to hydrogen sulfide-induced apoptosis between PHLDA1-overexpressing oral cancer cell lines and oral keratinocytes: Role of PHLDA1 as an apoptosis suppressor. Exp Cell Res [Internet]. 2014;320(2):247–57. Available from: http://dx.doi.org/10.1016/j.yexcr.2013.10.023

23. Sladky V, Schuler F, Fava LL, Villunger A. The resurrection of the PIDDosome - Emerging roles in the DNA-damage response and centrosome surveillance. J Cell Sci. 2017;130(22):3779–87.

24. Zhang X, Dong Y, Gao M, Hao M, Ren H, Guo L, et al. Knockdown of TRAP1 promotes cisplatin-induced apoptosis by promoting the ROS-dependent mitochondrial dysfunction in lung cancer cells. Mol Cell Biochem [Internet]. 2021;476(2):1075–82. Available from: https://doi.org/10.1007/s11010-020-03973-7

25. Zaccara S, Ries RJ, Jaffrey SR. Reading, writing and erasing mRNA methylation. Nat Rev Mol Cell Biol [Internet]. 2019;20(10):608–24. Available from: http://dx.doi.org/10.1038/s41580-019-0168-5

26. Chassé H, Boulben S, Costache V, Cormier P, Morales J. Analysis of translation using polysome profiling. Nucleic Acids Res. 2017;45(3):e15.

27. Oda E, Ohki R, Murasawa H, Nemoto J, Shibue T, Yamashita T, et al. Noxa, a BH3-Only Member of the Bcl-2 Family and Candidate Mediator of p53-Induced Apoptosis. 2000;288(May).

28. Xiang Y, Laurent B, Hsu CH, Nachtergaele S, Lu Z, Sheng W, et al. RNA m6 A methylation regulates the ultraviolet-induced DNA damage response. Nature. 2017;543(7646):573–6.

29. Shriwas O, Priyadarshini M, Samal SK, Rath R, Panda S, Majumdar SK Das, et al. DDX3 modulates cisplatin resistance in OSCC through ALKBH5-mediated m6A-demethylation of FOXM1 and NANOG. 2020.

30. Nie S, Zhang L, Liu J, Wan Y, Jiang Y, Yang J, et al. ALKBH5-HOXA10 loop-mediated JAK2 m6A demethylation and cisplatin resistance in epithelial ovarian cancer. J Exp Clin Cancer Res. 2021;40(1):1–18.

31. Tang F, Chen L, Gao H, Xiao D, Li X. m6A: An Emerging Role in Programmed Cell Death. Front Cell Dev Biol. 2022;10(January):1–9.

32. Meyer KD, Patil DP, Zhou J, Zinoviev A, Skabkin MA, Elemento O, et al. 5′ UTR m6A Promotes Cap-Independent Translation. Cell. 2015;163(4):999–1010.

33. Mao Y, Dong L, Liu XM, Guo J, Ma H, Shen B, et al. m6A in mRNA coding regions promotes translation via the RNA helicase-containing YTHDC2. Nat Commun [Internet]. 2019;10(1):1–11. Available from: http://dx.doi.org/10.1038/s41467-019-13317-9

34. Lin S, Choe J, Du P, Triboulet R, Gregory RI. The m6A Methyltransferase METTL3 Promotes Translation in Human Cancer Cells. Mol Cell [Internet]. 2016;62(3):335–45. Available from: http://dx.doi.org/10.1016/j.molcel.2016.03.021

35. Ploner C, Kofler R, Villunger A. Europe PMC Funders Group Noxa : at the tip of the balance between life and death. Eur PMC Funders Gr. 2012;27(Suppl 1):1–15.

36. Raj N, Wang M, Seoane JA, Moonie NA, Demeter J, Kaiser AM, et al. The Mettl3 epitranscriptomic writer amplifies p53 stress responses. 2021;

